# PrePR-CT: Predicting Perturbation Responses in Unseen Cell Types Using Cell-Type-Specific Graphs

**DOI:** 10.1101/2024.07.24.604816

**Authors:** Reem Alsulami, Robert Lehmann, Sumeer A. Khan, Vincenzo Lagani, David Gómez-Cabrero, Narsis A. Kiani, Jesper Tegner

## Abstract

Predicting the transcriptional response of chemical perturbations is crucial to understanding gene function and developing drug candidates, promising a streamlined drug development process. Single-cell sequencing has provided an ideal data basis for training machine learning models for this task. Recent advances in deep learning have led to significant improvements in predictions of chemical as well as genetic perturbations at the single cell level. Experiments have shown that different cell types exhibit distinct transcriptional patterns and responses to perturbation. This poses a fundamental problem for predicting transcriptional responses of drugs or cell types outside the training data. Accordingly, existing methods lack cell-type-specific modeling or do not explicitly provide an interpretable mechanism for the gene features. In this study, we introduce a novel approach that employs a network representation of various cell types as an inductive bias, improving prediction performance in scenarios with limited data while acknowledging cellular differences. We applied our framework to four small-scale single-cell perturbation datasets and one large-scale screening experiment, demonstrating that this representation can inherently generalize to previously unseen cell types. Furthermore, our method outperforms the state-of-the-art methods in predicting the post-perturbation response in unobserved cell types.

## 1 Introduction

In vitro drug screens, exemplified by High-Throughput Screening (HTS), enable the simultaneous testing of many molecules against specific biological activities or targets. Typically, HTS techniques are applied to bulk RNA sequencing data to quantify the transcriptional responses of thousands of molecules [1]. However, a major limitation of this approach is its disregard for cell-type-specific responses to individual perturbations. The transcriptional profiles generated by single-cell RNA sequencing methods provide a powerful resource for capturing the heterogeneity of cellular responses to various perturbations, including chemical compounds and CRISPR-based interventions. Nevertheless, the challenge of scaling HTS to single-cell resolution persists, primarily due to the substantial costs and technical complexities associated with highly multiplexed chemical experiments. Consequently, these experiments typically allow measuring the effects of a limited number of drugs, usually fewer than a hundred [2].

Recent experimental studies, such as [3, 4, 5, 6], have developed multiplexed single-cell transcriptional profiling techniques to investigate the heterogeneous responses of various cancer cell lines to chemical or genetic perturbations. These datasets inspire machine learning researchers to overcome experimental constraints by developing computational models capable of simulating single-cell gene expression profiles for unobserved perturbations. Many of these datasets are curated within the scPerturb project [2], which serves as a repository of standardized single-cell perturbation datasets.

Generative models, a category of deep learning methods, can generate new samples from the underlying data distribution. Existing models leverage generative approaches to map the data into a lower-dimensional latent space, subsequently employing simple vector arithmetic operations within this space to extrapolate the response of a specific drug on a different cell type. However, a prominent challenge with generative models is their demand for substantial data quantities to learn distribution parameters and generalize to novel conditions effectively. One of the solutions proposed by [7] involves applying transfer learning in conjunction with bulk RNA sequencing data, such as the connectivity map transcriptional data [1]. However, transfer learning restricts the feature space in the training set to the genes shared with the source domain dataset. This constraint may exclude crucial cell-type-specific marker genes and other contextually important features unique to certain cell types or conditions. Consequently, this limitation can reduce the model’s granularity and accuracy in predicting cell-type-specific responses, highlighting the need for methods that can integrate and preserve such cell-specific information. To address the limitations of transfer learning, recent studies have explored incorporating prior information, such as gene-gene interaction networks, to improve the prediction of cellular responses to novel perturbations [8]. While these models demonstrate that leveraging relational graphs can significantly enhance the accuracy of cellular response predictions, they still often overlook cell-type-specific differences between treatment and control groups. For instance a prior analysis by [9] compared these differences in the principal-component analysis (PCA) space and showed significant variability between the cell-type centroids in magnitude and direction. This analysis illustrates that the cell-type-specific context is crucial in predicting the perturbation responses.

To overcome the limitations of current generative models in predicting cell-type-specific responses to perturbations, we aim to leverage prior information, specifically cell-type-specific co-expression networks, to predict the mean and variance of post-perturbation expression levels for Differentially Expressed Genes (DEGs) in previously unobserved cell types. This study systematically evaluates the performance of our proposed method across four small-scale single-cell perturbation datasets and one large-scale chemical screening experiment. Additionally, we conduct a comparative analysis with two state-of-the-art generative models, focusing exclusively on chemical perturbations to benchmark our approach. Our results demonstrate the effectiveness of our framework in addressing the challenges associated with predicting DEGs expression in diverse cellular contexts. The proposed framework employs the graph attention network (GAT) algorithm [10] to extract salient features, enabling a more accurate representation of cell-type-specific gene interactions. To investigate the interpretability of this approach, we analyze the attention values learned by the GAT layers for each gene, revealing their capability to capture non-trivial patterns from the data.

## 2 Method

Our method considers a perturbational single-cell dataset 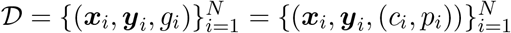,where ***x***_*i*_ and ***y***_*i*_ ∈ ℝ^*m*^ are m-dimensional gene expression vectors representing a control cell and a cellular response to a group of perturbation *g*_*i*_, respectively. Each group *g*_*i*_ consists of a cell type *c*_*i*_ ∈ 𝒞 and a *k*-dimensional learnable embedding vector *p*_*i*_ representing a perturbation. We associate each cell type with one of the cell-type-specific graphs 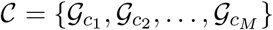, where *M* is the number of cell types in 𝒟. We predict the perturbation effect as a function of the cell type graph, a set of pre-selected control samples of the same cell type, and a perturbation embedding.

### 2.1 Cell-type-specific graphs

Cell-type specific gene expression profiles of cells in an unperturbed state are modeled using co-expression graphs, which are constructed as follows. Single-cell expression values of unperturbed cells are aggregated into metacells using SEACells [11] to mitigate the inherent sparsity of single-cell expression data. The set of 5000 Highly Variable Genes (HVG) is then calculated across all metacells. To construct the graph for a given cell type 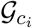, all pairwise correlations between HVGs is calculated using only metacells of the selected cell type. Weaker correlating pairs are then filtered out using the 99th percentile of the absolute correlations in the selected cell type as the threshold. Since we assume no access to perturbed expression data at testing time, corresponding graphs can not be constructed for perturbations.

The resulting co-expression graphs include varying numbers of genes based on the defined threshold, with some genes shared across cell types. These shared genes can be used as a similarity measure between the cell type graphs. The selected parameters depend on the processed data and constitute a trade-off between considering the most specific genes per cell type, retaining similar genes between the cell type graphs, and controlling the graph’s size, which affects the time complexity of the training.

The feature matrix of the graph is initialized with *k*-dimensional learnable embedding vectors representing the genes, 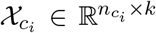, where 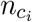 is the number of genes in 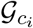. These embeddings are defined so that a given gene has the same representation across the cell graphs. This enables the model to be aware of the overlapped genes that preserve the similarity between the cell types. Additionally, they are updated during the backpropagation step in the training process to minimize the loss function. These gene embeddings are combined with the mean and standard deviation of the control cells in the feature matrix to obtain the basal expression state.

After constructing the cell-type-specific graphs 𝒞, we measure cell-type similarity using the number of shared genes between the representative graphs. Using the resulting similarity matrix, we identify the cell type that is the most dissimilar from the majority of the remaining cell types and reserve it for testing.

### 2.2 Matching control and perturbed cells

One challenge in predicting transcriptional responses is the lack of paired measurements in the same cell before and after perturbation due to the destructive nature of single-cell transcriptomic measurement techniques. Furthermore, in the considered datasets there tend to be more cells available in the control state compared to the perturbed state, requiring subsampling of control state cells. We employ the SEACells metacell clusters, which are used to construct co-expression graphs, to ensure an even sampling of control cells across the available dataset independent of cell density. For each cell type and perturbation, we evenly sample individual cells from the control state metacell clusters to match the number of perturbed cells, which reduces the number of samples in the data and thus the time complexity of the training process. It is important to note that we are using the gene expression samples as both input and output; the SEACells clusters serve merely as a tool to align the two groups.

### 2.3 Chemical Structure Representation of Perturbing Compound

We utilize SMILES (Simplified Molecular Input Line Entry System) chemical structures to represent the chemical perturbations under investigation. SMILES notation offers a compact and machine-readable approach to describing molecular structures. These representations serve as the input for generating molecular embeddings using the ChemBERTa model [12], which employs state-of-the-art natural language processing techniques adapted specifically for chemical data. ChemBERTa was pre-trained using a dataset containing 77 million distinct SMILES obtained from PubChem [13], recognized as the largest publicly accessible repository of chemical structures worldwide. It leverages transfer learning from the RoBERTa transformer to learn rich representations of molecular structures and their associated properties [12].

This approach enables us to capture nuanced structural features and chemical properties encoded within molecular structures. We obtained the SMILES representations of the perturbations used in this study from the DrugBank database [14] or from the original publication of the corresponding dataset. By leveraging ChemBERTa embeddings, our model integrates both molecular structure information and transcriptional response data to predict drug effects across different cell types and conditions.

### 2.4 Graph Attention Network

The Graph Attention Network (GAT) defines an attention mechanism on the graph structure by computing an attention score *α*_*i,j*_ between a given node *i* and each node *j* in its first-order neighborhood set 𝒩_*i*_ [10]. This score represents the significance of node *j*’s features to node *i* and is calculated as:

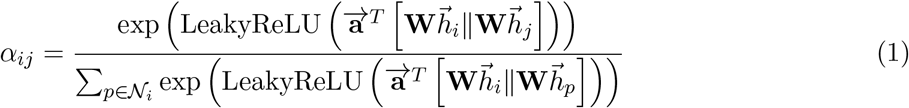

where 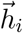 is a feature representation of node *i* transformed using the weight matrix **W**, and **a** is a learnable weight vector. The self-attention mechanism is then extended to multi-head attention by applying *L* independent mechanisms, and their outputs are either concatenated or averaged:

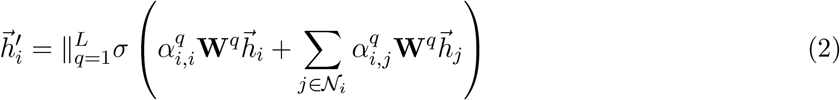

The final output 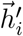 is a high-level feature representation of node *i* in the graph. We employed GAT in this project because it has shown successful examples of generalizing to completely unobserved graphs during the training process [10]. See step 3 in figure 1.

**Figure 1:**
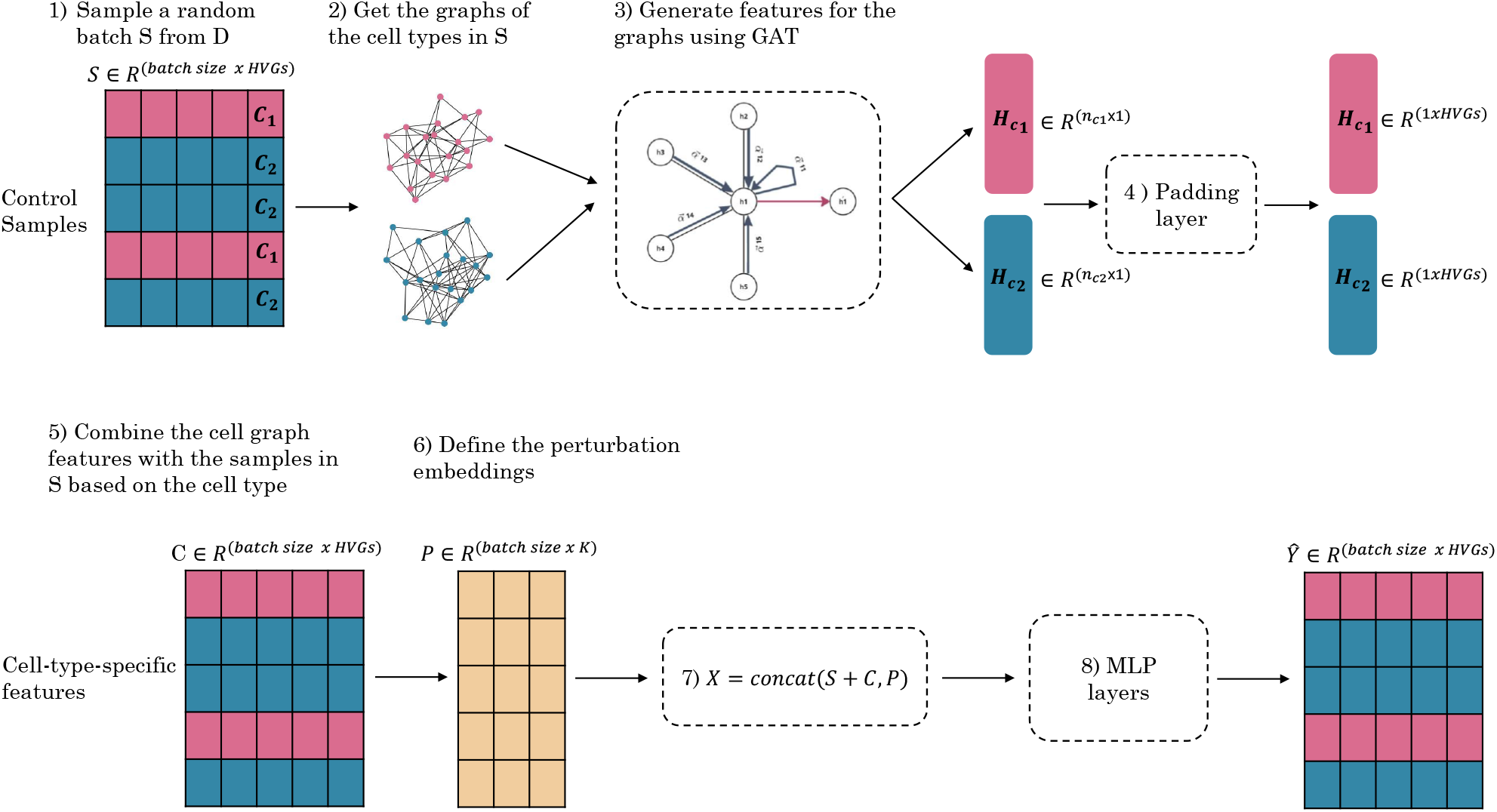
Training steps of PrePR-CT. PrePR-CT encodes cell-type graphs from a batch of training samples using GAT layers, integrates them with control gene expression and pre-defined perturbation embeddings, and processes them through Multi-perceptron layers to predict the gene expression response. The different colors in the rows of the matrices represent the cell type labels.

### 2.5 Loss Function

The Mean Square Error (MSE) is commonly used as a loss function in the training of regression models. However, MSE primarily focuses on the mean and does not take into account the overall distribution of the actual and predicted samples. In contrast, the Wasserstein distance, also known as the Earth Mover’s Distance (EMD), is an optimal transport (OT) measure for comparing probability distributions with disjoint supports.[15] The Wasserstein distance has achieved significant success in the context of Generative Adversarial Networks (GANs). The introduction of the Wasserstein GAN (WGAN) framework has led to more stable training and improved the quality of generated samples by providing a more meaningful and smooth gradient signal compared to traditional GANs [16]. To approximate the Earth Mover’s Distance, we seek the optimal transportation cost *γ*:

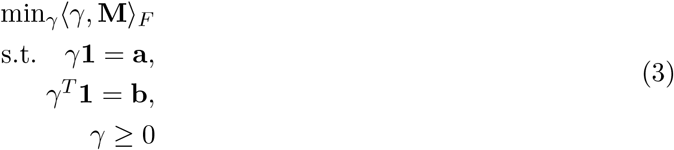

where **M** is a cost matrix corresponding to the Euclidean distances between the actual and predicted samples, and **a** and **b** are the sample weights of the source and target distributions, respectively [17, 18]. For the implementation, we used the minibatch EMD function implemented by [19] to solve the aforementioned optimization problem. The optimal transport library in Python offers several tutorials for using this loss function [19].

The final loss is defined as:

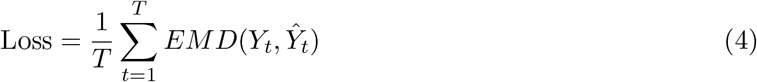

Where *T* represents the number of cell type and perturbation pairs *g*_*i*_ in a given batch of samples, and *Y*_*t*_ as well as *Ŷ*_*t*_ are the actual and predicted samples of *g*_*i*_ in the current training batch, respectively.

### 2.6 Implementation details

Utilizing a cell-type-specific gene interaction network requires mapping each gene expression sample to the graph of its respective cell type. However, this approach increases the time complexity of the training process. To address this issue, we adopt an alternative formulation. First, we extract the graphs within a given batch of samples and then embed them using graph attention layers. Then, we map the samples to the features generated using the GAT layers based on the cell type labels. This approach prevents us from treating each cell as a separate graph, thus reducing computational complexity. The proposed method incorporates the following inputs: 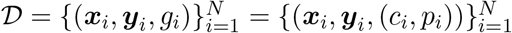 and 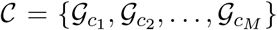. For details, refer to the Method section 2. Figure 1 demonstrates the training steps.

The model required tuning six hyperparameters during training. Table 1 lists these parameters along with the values optimized throughout the training process using the hyperparameter optimization tool Optuna [20] on a fixed random validation set. We employed the Adam optimizer [21] for model optimization, and no activation function was applied to the output layer.

**Table 1:**
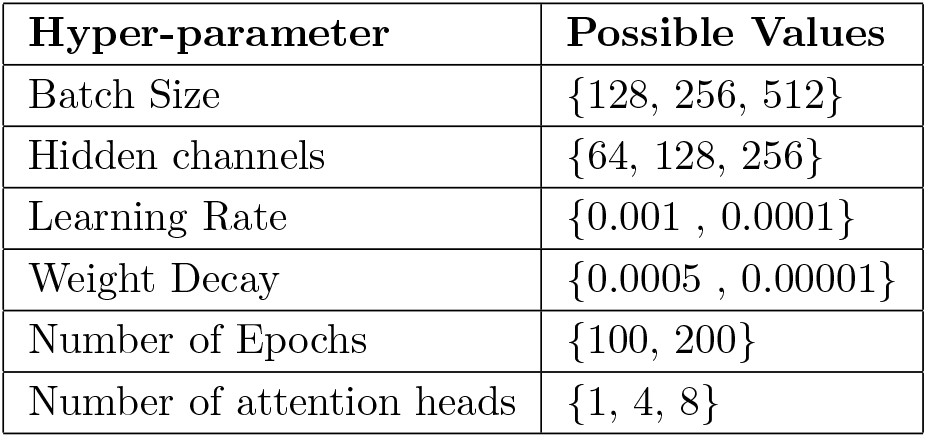
Overview of the hyper-parameters investigated for each dataset in this study.

| Hyper-parameter | Possible Values |
| --- | --- |
| Batch Size | {128, 256, 512} |
| Hidden channels | {64, 128, 256} |
| Learning Rate | {0.001 , 0.0001} |
| Weight Decay | {0.0005 , 0.00001} |
| Number of Epochs | {100, 200} |
| Number of attention heads | {1, 4, 8} |

**Table 2:** Overview of the datasets used in this study.

| Dataset | Source | Control Samples | Perturbed Samples | Cell Types | Perturbations |
| --- | --- | --- | --- | --- | --- |
| Kang | GEO - GSE96583 [22] | 12,388 | 11,861 | 7 | 1 |
| NeurIPS | Kaggle [24] | 28,461 | 56,668 | 6 | 6 |
| McFarland | scPerturb Project [2] | 2,917 | 13,291 | 5 | 11 |
| Nault | GEO - GSE184506 [23] | 12,452 | 6,282 | 6 | 1 |
| Chang | scPerturb Project [2] | 21,006 | 21,230 | 1 | 3 |

### 2.7 Evaluation

We employ the coefficient of determination (*R*^2^) to assess the alignment between observed data statistics (mean and standard deviation) and the corresponding prediction. Our analysis specifically focuses on gene expression statistics rather than overall gene expression profiles. This distinction is crucial because other methods included in the comparative analysis map all control cells to the perturbed state, resulting in an unequal number of output samples compared to the actual perturbed samples in the ground truth data.

Consequently, we concentrate on capturing the distinct shifts from the control state to the perturbed state. The intentional choice of *R*^2^ as our evaluation metric offers notable advantages over alternative metrics such as Pearson correlation and Mean Squared Error (MSE). Unlike Pearson correlation, which measures only linear relationships and overlooks magnitude considerations, *R*^2^ provides a more comprehensive assessment of predictive power. This deliberate selection aligns with our objective of capturing nuanced changes in gene expression.

For the gene set selected for evaluation, we assess all the genes in the graph of the testing cell type and the top 100 DEGs. The top 100 DEGs are defined using a combined ranking scheme after computing the p-value and log fold change using t-tests in the `scanpy` package in Python. First, we filter genes with a p-value of 0.05 and then apply the rank product statistic:

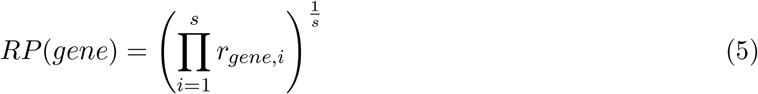

where *s = 2 corresponds to the* p-value and log fold change. The rationale behind this ranking is to eliminate noisy genes that have extreme log fold change values and very small p-values.

## 3 Datasets

We assess the efficacy and versatility of our method across diverse scenarios by evaluating it on multiple datasets encompassing various cell types and perturbations. Specifically, we employ five distinct datasets: human peripheral blood mononuclear cells (PBMCs) treated with interferon *β (IFN-β) publishe*d by Kang et al.[22], McFarland dataset [3], Chang dataset [4], single-dose TCDD liver by Nault et al.[23], and a subset of the cross-individual dataset (NeurIPS) from the Open Problems in Single-cell competition [24].

The McFarland dataset encompasses 13 chemical perturbations, each with a specific dose value, and comprises over 100 cell lines. In our analysis, we narrowed our focus to the top five cell lines based on sample count and excluded drugs lacking SMILES identifiers in the DrugBank database [14].

The Nault dataset entailed single-nucleus RNA sequencing (snRNA-seq) of flash-frozen C57BL6 mouse livers. In this dataset, mice underwent subchronic administration of a specified TCDD dose via oral gavage every four days over a span of 28 days. As indicated by [9], we excluded all immune cell types, given their propensity to migrate from the lymph to the liver during TCDD administration.

The NeurIPS dataset comprises single-cell gene expression profiles obtained from human peripheral blood mononuclear cells (PBMCs) treated with 144 compounds sourced from the LINCS Connectivity Map database [1]. Measurements were conducted 24 hours post-treatment, with the experiment replicated using cells from three healthy human donors. To streamline our analysis, we selected a subset of perturbations based on the median absolute log-fold change across all genes and cell types. This process led to the identification of six drugs exhibiting the most pronounced responses. It is important to note that only 15% of the perturbations are available in the Meyloid and B cell types. Consequently, our subset was chosen from these specific perturbations.

We processed all the datasets through a series of steps, which involved removing mitochondrial and ribosomal genes, filtering for cells with a minimum of 1000 counts and genes detected in fewer than 50 cells, normalizing the count matrix using the `normalize_total` function in the `scanpy.pp` package in Python, and applying a log(*x* + 1) transformation.

For the cell type graphs, we followed the procedure described in Section 2 to construct the graphs for each dataset. The total gene-gene correlation matrix contains 5000 × 5000 edges. By applying thresholding, we retain between 240,000 and 250,000 edges per cell type. The number of nodes in each graph is provided in Section 4, with examples shown in Figure 2.

**Figure 2:**
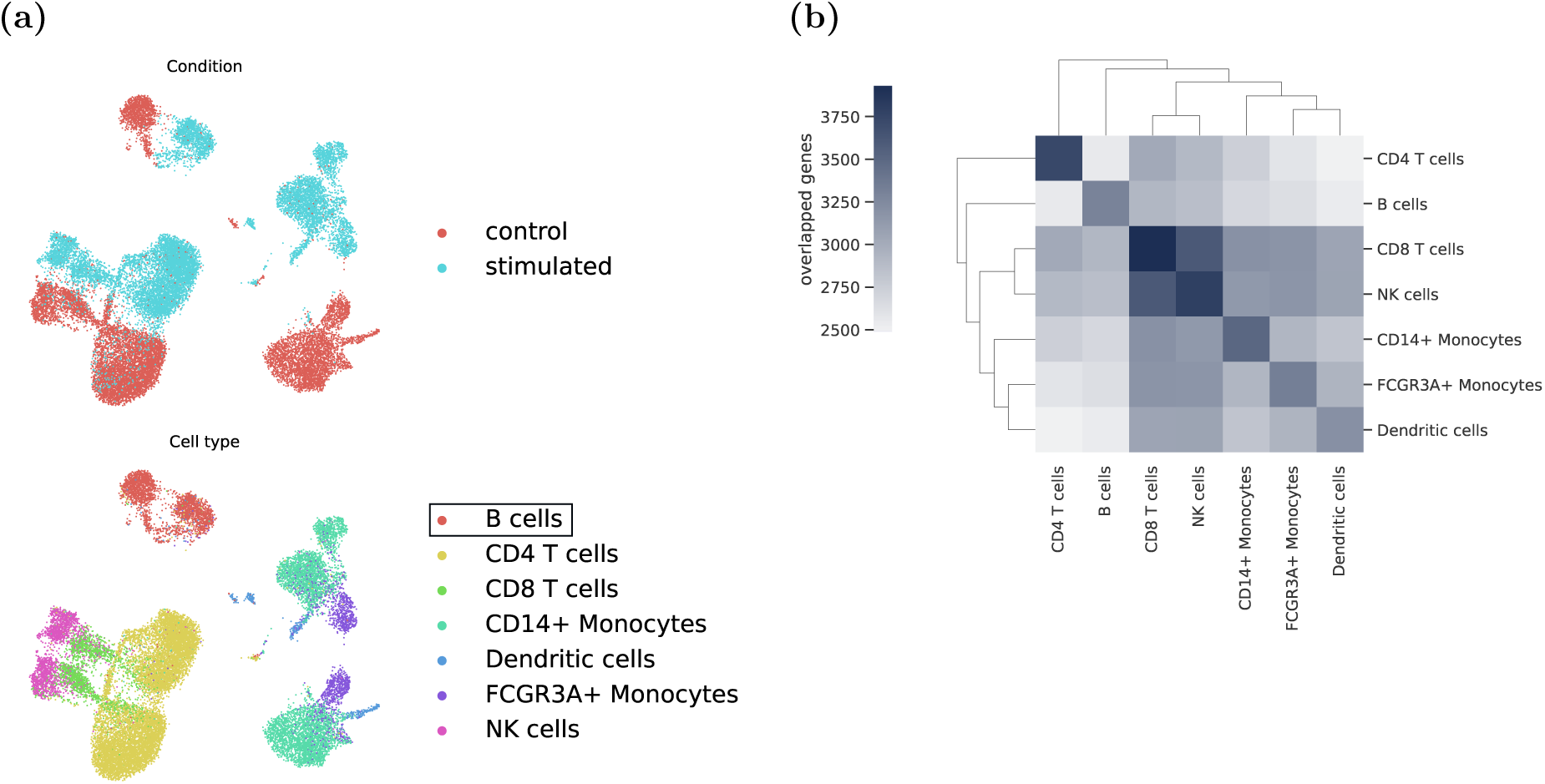
Single-perturbation experiment with Kang dataset. (a) UMAP Projection showing the testing cell type stimulated with IFN-*β*. (b) A clustered heatmap showing the number of shared genes/nodes between the cell type graph pairs.

## 4 Experiments and Results

PrePR-CT is a graph-based deep learning method designed to predict transcriptional responses to chemical perturbations in single-cell data using cell-type-specific co-expression networks. The method employs GAT layers to encode cell-type graphs from batches of training samples. These encoded graphs are then integrated with control gene expression data and predefined perturbation embeddings. The combined data is processed through Multi-Layer Perceptrons (MLPs) to accurately predict gene expression responses. In this section, we present the results and analysis for both scenarios. We showcase a titration experiment on the Kang dataset to demonstrate our model’s capability in a small data regime. Additionally, we perform a comparative analysis with a state-of-the-art method for predicting perturbation responses in unseen cell types. Finally, we analyze the high attention genes (HAGs) in the Kang dataset identified through the GAT layers, demonstrating their ability to extract non-trivial features from the data.

### 4.1 Cell-type-specific graphs preserve the similarities between the cell types

Before evaluating our model’s ability to predict perturbation responses, we first demonstrate the reliability of our graph representation in accurately capturing the inherent data structure. This ensures the preservation of similarities between cell types, as indicated by the distances between their clusters in the UMAP space. This observation allows us to use it as a selection criterion for the testing cell type, where different cell types in the UMAP space exhibit distinct gene patterns in the cell type graphs. The quantification of these similarities is achieved by computing the number of overlapped genes between the cell type graphs. Consequently, the model’s ability to generalize to unseen cell types stems from leveraging interactions with these shared genes among the cell type graphs. To validate this observation, we construct cell-type-specific graphs for four distinct single-cell perturbation datasets using the procedure outlined in section 2.

The Kang dataset consists of samples from seven distinct cell types exposed to INF-*β*. Figure 2a illustrates the UMAP embedding of the data, labeled by both condition and cell types, while Figure 2b depicts the corresponding overlap matrix derived from the cell type graphs. Notably, as observed in the UMAP figure, the node sets of CD8 T cells and NK cells exhibit a high degree of similarity, whereas the B cells cluster appears notably distant from the rest of the cell types. We designate this cell type for testing, assuming it to be challenging for the model to predict its response compared to the other cell types.

Additionally, we examine the similarity patterns among the PBMCs in the NeurIPS dataset, as illustrated in Figure 3. We observe clustering between T cells CD8+ and T regulatory cells, as well as between T cells CD4+ and NK cells. Similar to the Kang dataset, the B cells cluster is the most distant from the other cell types. Additional datasets are provided in the supplementary material (see Appendix 6).

**Figure 3:**
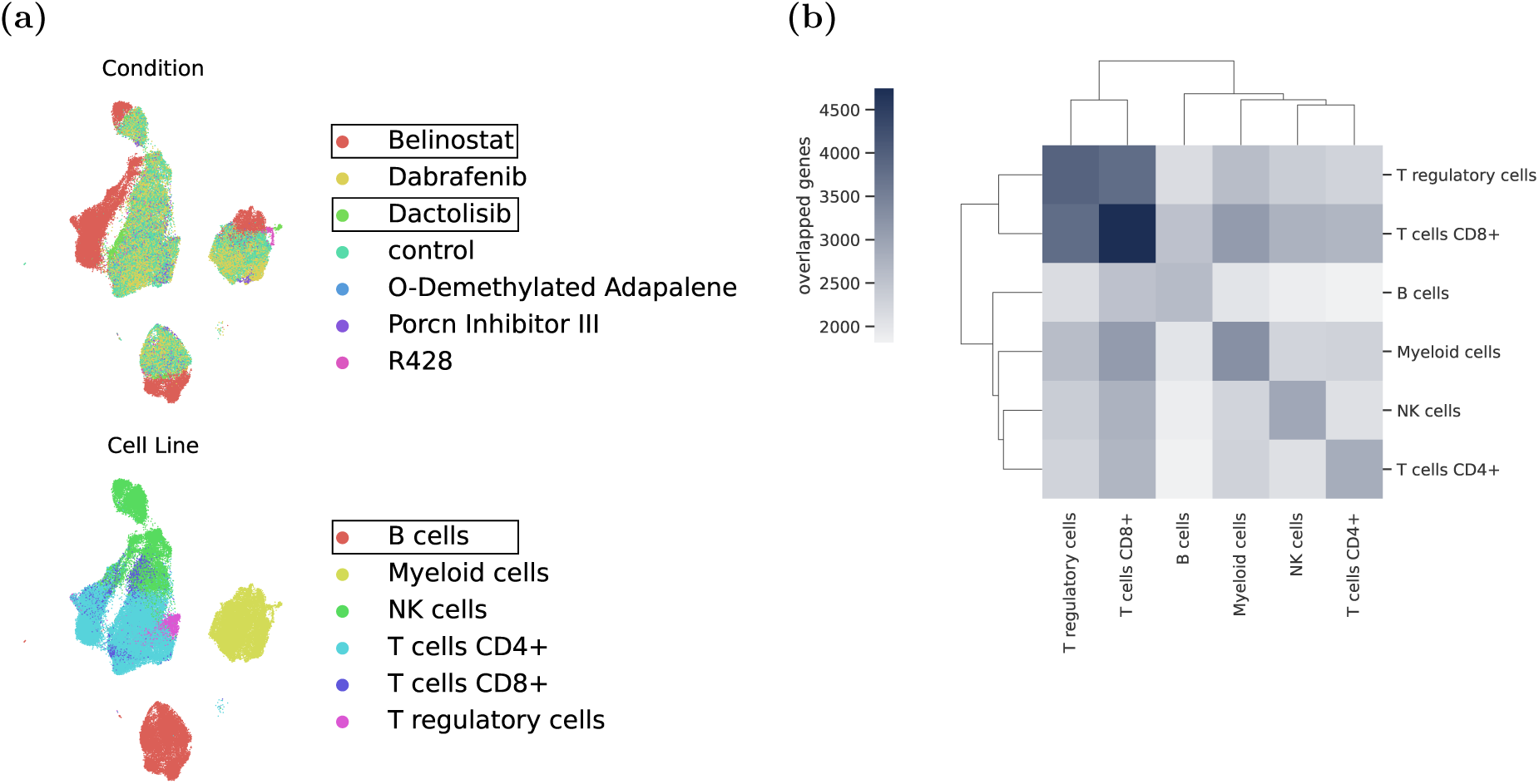
Multi-perturbation experiment with NeurIPS dataset. (a) UMAP Projection showing the testing cell type and drugs (b) A clustered heatmap showing the number of shared genes/nodes between the cell type graph pairs.

### 4.2 Predicting the perturbation response in unseen cell type

After generating cell-type-specific graphs for each dataset and identifying the target cell type for testing, we apply the E-distance metric proposed by Peidli et al. [2] to identify drugs that exhibit the most significant deviation from the control cells within the selected testing cell type. These drugs are then selected for testing as they represent the most challenging perturbations to predict. Subsequently, we train PrePR-CT using different datasets. The training procedure follows the methodology outlined in Section 1. Given that our approach involves input pairs of cell types and drugs, we compare its performance with established baselines such as scGen [25], CPA on single-perturbation datasets [26], and chemCPA [7] on multi-perturbation datasets, utilizing molecular structures as features for the perturbations (see Section 2.3).

First, we consider scenarios involving multiple cell types to predict the perturbation response of unseen cell types. We evaluate our method using four datasets: Kang, Nault, McFarland, and NeurIPS. Figures 4 and 5 demonstrate its performance in predicting the mean and variance of different perturbations. We use different colors to distinguish the top 20 DEGs, dividing them into red points and green points. Red points indicate up-regulated genes (increased compared to the control), while green points represent down-regulated genes. Notably, PreRP-CT performs well in predicting the top 100 DEGs. Additionally, the correlation plots demonstrate the performance in predicting the response to different perturbations, indicating that some perturbations down-regulate the gene expression while others increase it.

**Figure 4:**
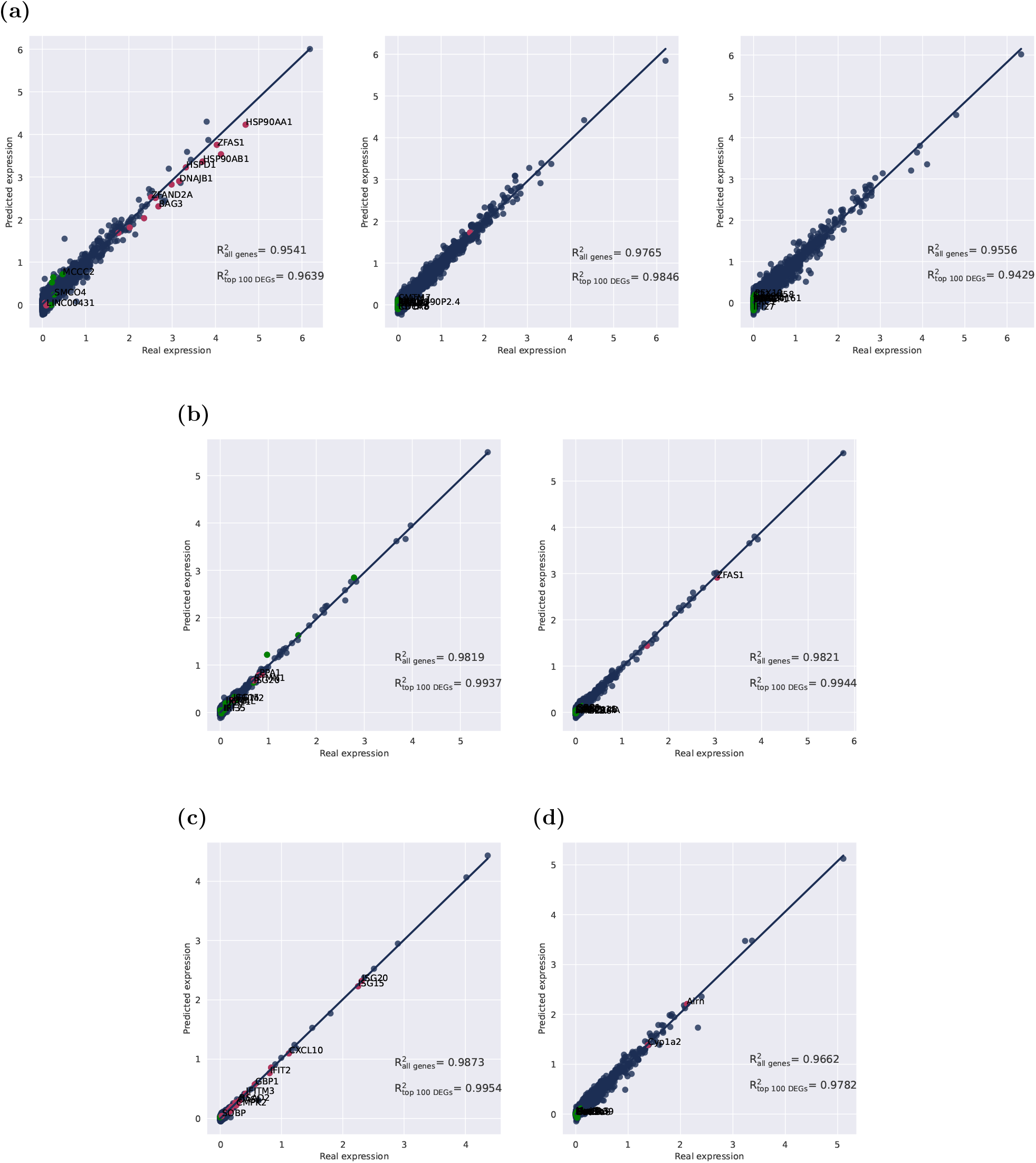
Correlation between mean gene expression values of actual and predicted responses. (A) McFarland (Bortezomib, Gemcitabine, and JQ1) drugs. (B) NeurIPS (Belinostat and Dactolisib). (C) Kang (IFN-*β*). (D) Nault (TCDD).

**Figure 5:**
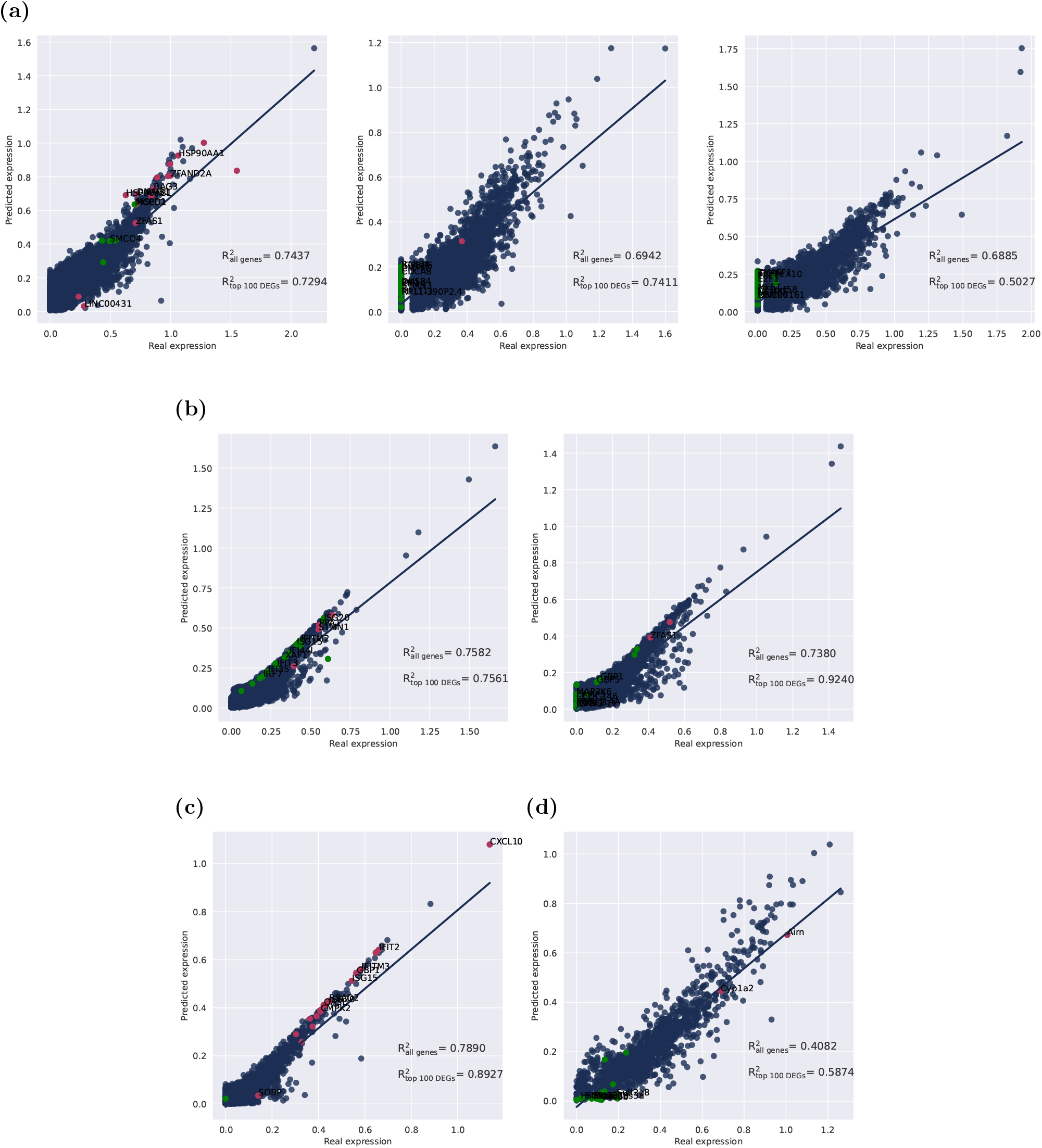
Correlation between standard deviations of gene expression values for actual and predicted responses. (A) McFarland (Bortezomib, Gemcitabine, and JQ1) drugs. (B) NeurIPS (Belinostat and Dactolisib). (C) Kang (IFN-*β*). (D) Nault (TCDD).

Figure 4 compares the observed and predicted mean gene expression across various datasets, while Figure 5 compares the standard deviations to evaluate the variability prediction task.

Besides the correlation plots, we show a UMAP projection of the actual and predicted gene expression of the testing cell type and drugs against the control cells in Figure 6. The figure reveals a significant overlap between the observed and predicted cells in the UMAP projection, underscoring the effectiveness of our method in capturing the transcriptional response to unseen perturbations.

**Figure 6:**
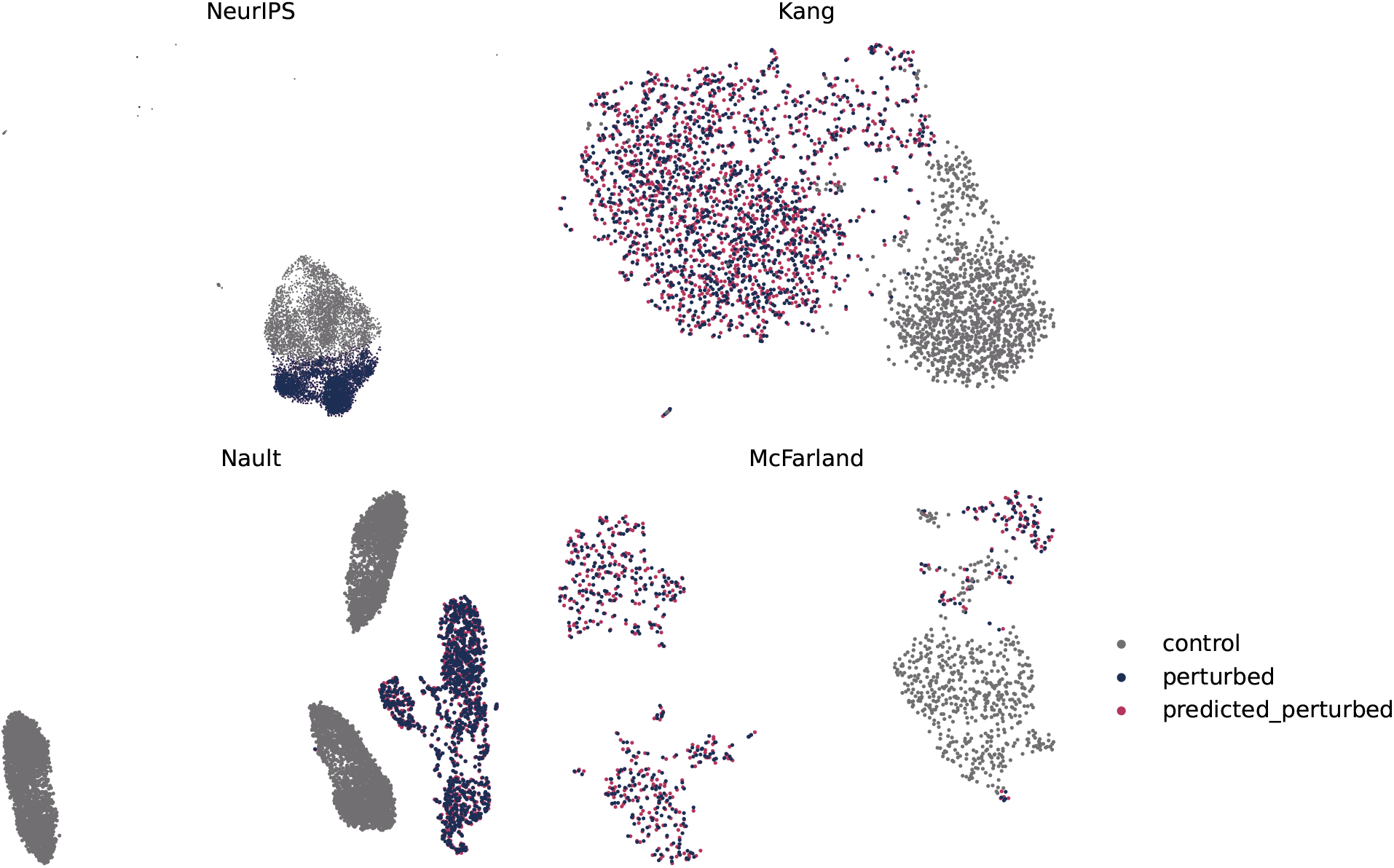
UMAP projection comparing the observed and predicted gene expression.

As previously mentioned, we conducted comparisons with two state-of-the-art generative models in predicting the effects of perturbations. scGen faces limitations in scenarios with only one cell line and multiple drugs, as each drug must be represented in the training set with different cell types. In light of this, we followed the steps outlined in the official documentation of scGen and CPA to train and test the models (https://scgen.readthedocs.io/en/stable/tutorials/scgen_perturbation_prediction.html, https://cpa-tools.readthedocs.io/en/latest/tutorials/Kang.html). All models underwent comparison using the same testing cell types and drugs.

We trained scGen and CPA on the top 5000 highly variable genes, which were also used to construct the cell type graphs. During testing, evaluation was performed on all genes within the graph of the testing cell type, specifically focusing on the top 100 DEGs. Tables 3 and 4 below compare the performance of the three methods across multiple datasets. For the multiple-perturbation datasets, we considered the average *R*^2^ of the testing drugs. The results indicate that PrePR-CT significantly outperformed scGen and CPA/chemCPA, especially in variability prediction. Also, we compare the observed and predicted gene expression states of the top 20 DEGs, as shown in Figure 7. These comparisons demonstrate that PrePR-CT consistently provides more accurate predictions of gene expression states than scGen and CPA/chemCPA, highlighting its performance in capturing the complexity of gene expression variability.

**Table 3:** Comparison of our method with the scGen and CPA models for predicting the mean post perturbation responses in unseen cell types across four datasets.

| Dataset | Testing cell type | Testing drug | Gene Set | CPA | scGen | PrePR-CT |
| --- | --- | --- | --- | --- | --- | --- |
| Kang | B cells | IFN-β | All Genes | 0.9433 | 0.9663 | **0.9873** |
|  |  |  | 100 DEGs | 0.9592 | 0.9537 | **0.9954** |
| Nault | Hepatocytes - portal | Nault | All Genes | 0.6271 | 0.8096 | **0.9662** |
|  |  |  | 100 DEGs | 0.7674 | 0.6251 | **0.9782** |
| McFarland | LNCAPCLONEFGC | JQ1, Bortezomib<br>Gemcitabine | All Genes | 0.8404 | 0.8244 | **0.9621** |
|  |  |  | 100 DEGs | 0.7728 | 0.4254 | **0.9637** |
| NeurIPS | B cells | Belinostat, Dactolisib | All Genes | 0.9511 | 0.9558 | **0.9820** |
|  |  |  | 100 DEGs | 0.9717 | 0.92416 | **0.9940** |

**Table 4:** Comparison of our method with the scGen and CPA models for predicting the standard deviation of post-perturbation responses in unseen cell types across four datasets.

| Dataset | Testing cell type | Testing drug | Gene Set | CPA | scGen | PrePR-CT |
| --- | --- | --- | --- | --- | --- | --- |
| Kang | B cells | IFN-β | All Genes | 0.2665 | -0.2563 | **0.7890** |
|  |  |  | 100 DEGs | 0.2945 | -0.4020 | **0.8927** |
| Nault | Hepatocytes - portal | Nault | All Genes | -0.0311 | -1.4786 | **0.4082** |
|  |  |  | 100 DEGs | 0.3007 | -0.1640 | **0.5874** |
| McFarland | LNCAPCLONEFGC | JQ1, Bortezomib<br>Gemcitabine | All Genes | **0.7215** | 0.1209 | 0.7087 |
|  |  |  | 100 DEGs | **0.7309** | 0.3591 | 0.6577 |
| NeurIPS | B cells | Belinostat, Dactolisib | All Genes | 0.1966 | -0.3320 | **0.7480** |
|  |  |  | 100 DEGs | 0.4469 | 0.3800 | **0.8400** |

**Figure 7:**
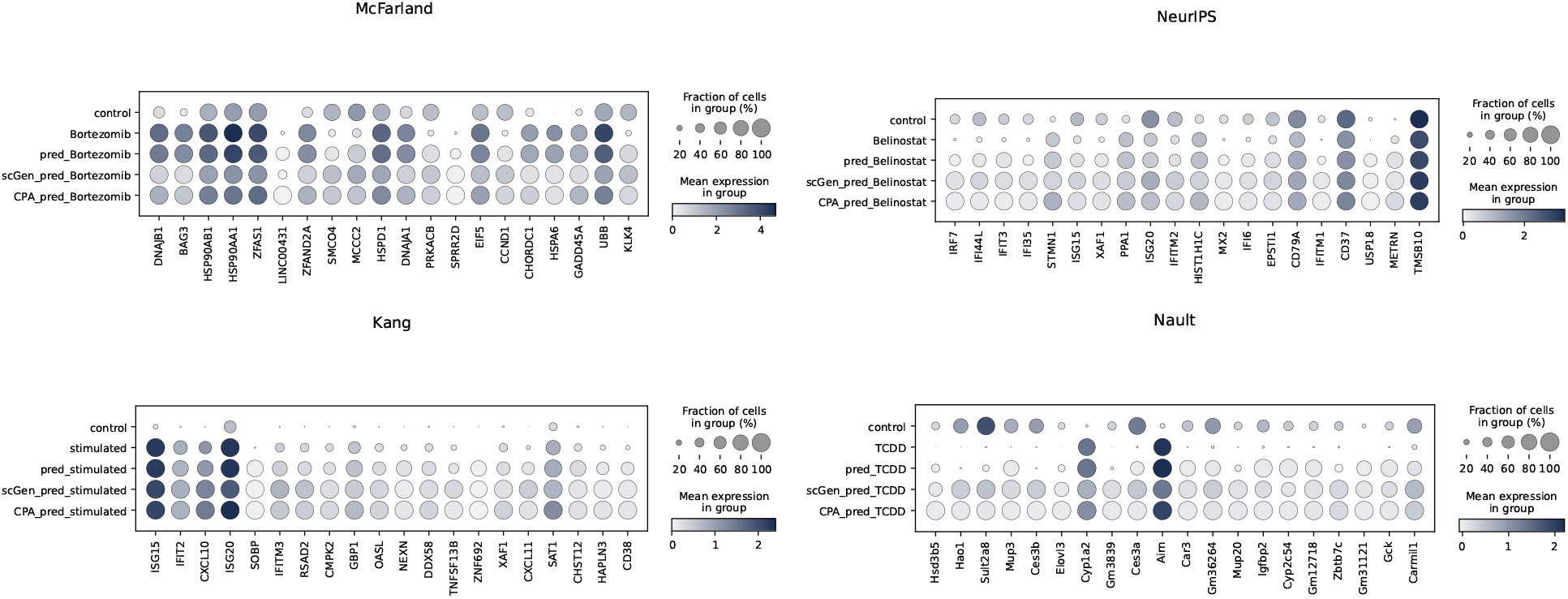
Dot plot comparing the actual and predicted expression of the top 20 DEGs.

### 4.3 Predicting the response of unseen perturbation in the same cell type

In addition to predicting the perturbation effect in unseen cell types, we also explore the scenario of predicting the response of previously unseen drugs within the same cell type. For this purpose, we utilize the Chang dataset [4]. This dataset exclusively includes the lung cancer cell line PC9, which was exposed and treated for four days with three structurally similar drugs: 1 μM erlotinib, 1 μM GNE-069, and 1 μM GNE-104, respectively. The UMAP visualization reveals a significant separation between control and treated cells while demonstrating similar responses among the drugs. GNE-069 was excluded from the training set based on its Euclidean distance from the control cells. The testing performance in Figure 8 showcases the potential of our method to predict unseen perturbation effects, even in datasets with a limited number of perturbations. Since scGen cannot predict the effect of unseen perturbations, we conducted a comparison with the chemCPA model. The results indicate that our method significantly outperforms chemCPA across different gene sets, as shown in Tables 5 and 6.

**Table 5:** Comparison of our method with the CPA model for predicting the mean response of unseen perturbation in the same cell type in Chang dataset.

| Dataset | Testing cell type | Testing drug | Gene Set | CPA | PrePR-CT |
| --- | --- | --- | --- | --- | --- |
| Chang | PC9 | GNE-069 | All Genes | 0.9825 | **0.9949** |
|  |  |  | 100 DEGs | 0.9869 | **0.9955** |
|  |  |  | 50 DEGs | 0.9928 | **0.9975** |
|  |  |  | 20 DEGs | 0.9909 | **0.9930** |

**Table 6:** Comparison of our method with the CPA model for predicting the standard deviation of the response of unseen perturbation in the same cell type in Chang dataset.

| Dataset | Testing cell type | Testing drug | Gene Set | CPA | PrePR-CT |
| --- | --- | --- | --- | --- | --- |
| Chang | PC9 | GNE-069 | All Genes | 0.2708 | **0.5447** |
|  |  |  | 100 DEGs | 0.8122 | **0.8985** |
|  |  |  | 50 DEGs | 0.8810 | **0.9364** |
|  |  |  | 20 DEGs | 0.92578 | **0.9660** |

**Figure 8:**
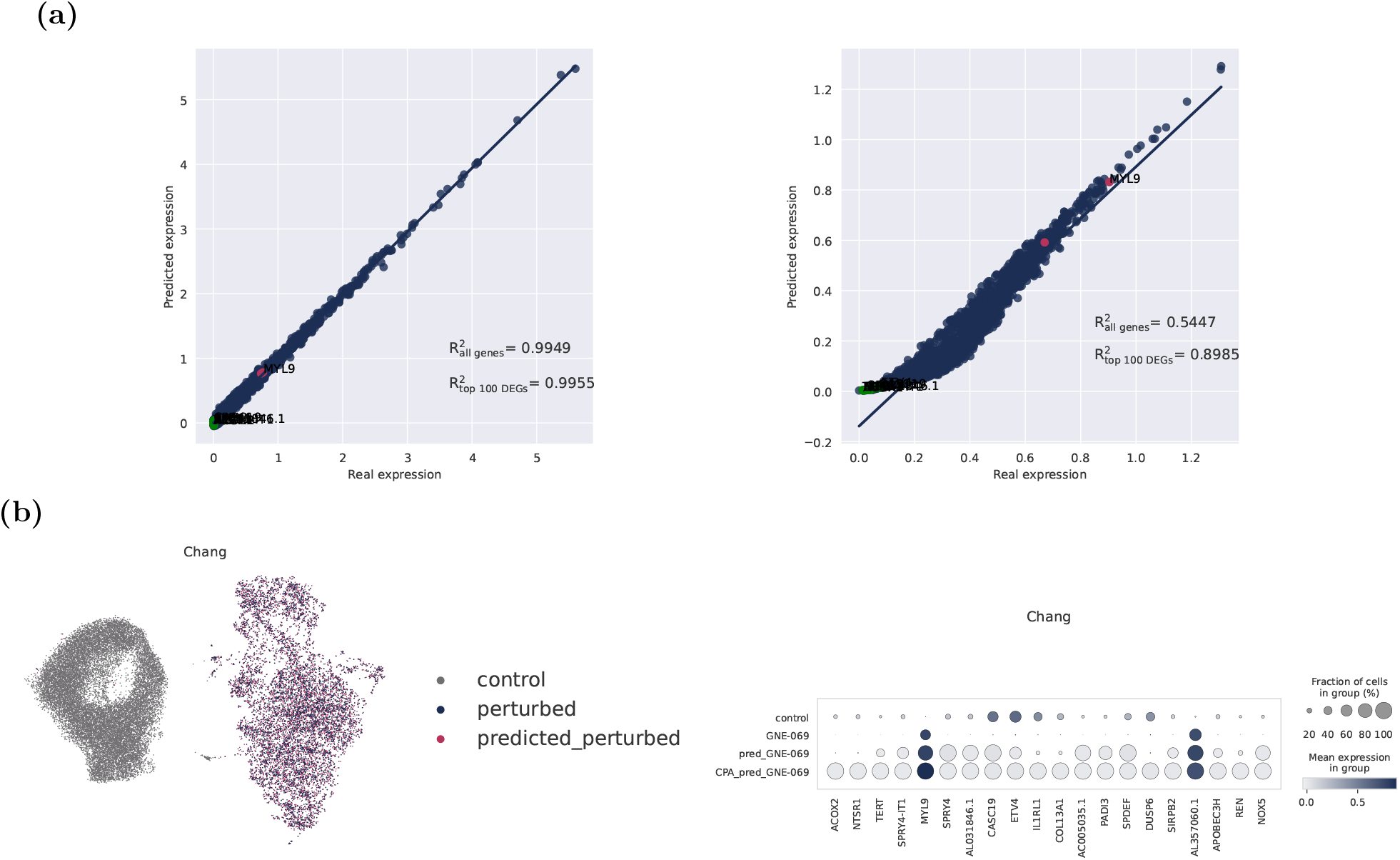
Results of predicting the unseen perturbation (GNE-069) in the Change dataset. (a) Correlation plots comparing the mean and standard deviation. (b) UMAP projection to assess the alignment between observed and predicted gene expression. (c) Comparison of gene expression states for the top 20 DEGs.

### 4.4 Predicting the perturbation response in small data regime

In the preceding section, we showcased the performance of PrePR-CT in predicting post-perturbation responses using different datasets. In this section, our focus shifts to a more comprehensive investigation. We trained our model on subsets derived from the Kang dataset, encompassing varying sample sizes of the entire dataset, including both perturbed and control samples.

Figure 9 presents the *R*^2^ values for PrePR-CT and CPA methods evaluated across all genes and the top 100 DEGs at different sample sizes. The figure illustrates a consistent enhancement in PrePR-CT’s performance as the sample size increases, indicating sensitivity of prediction validity to sample size. Importantly, our model’s effectiveness extends beyond larger datasets; even in scenarios with smaller subsets, it consistently demonstrates commendable performance. From 60% of the dataset up to the entire dataset, the model shows nearly constant performance. Furthermore, we observe nuanced distinctions between scenarios involving all genes and those with only 100 DEGs. One plausible explanation is that the top DEGs represent genes showing the maximum change, thereby effectively distinguishing between treatment and control groups.

**Figure 9:**
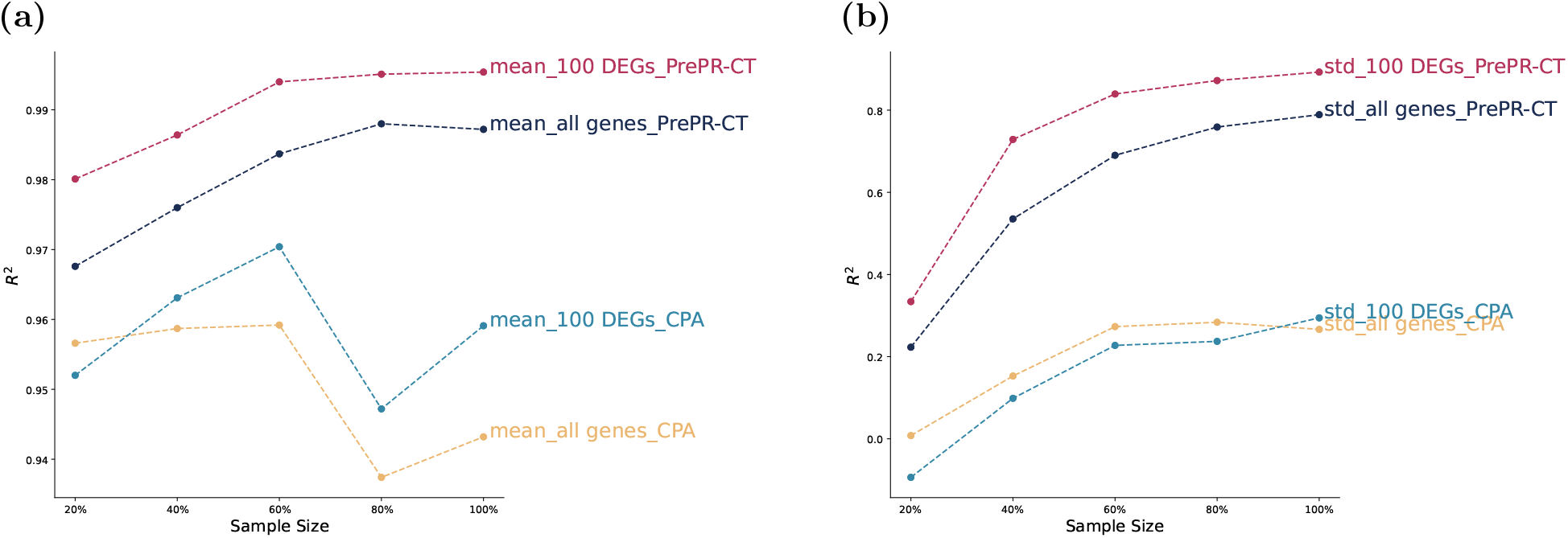
The Coefficient of Determination (*R*^2^) metric comparison between PrePR-CT and CPA across different fractions of training samples within the Kang dataset. Yellow and light blue lines represent CPA, while the remaining lines depict PrePR-CT. (a) Prediction of mean post-perturbation gene expression. (b) Prediction of the standard deviation in post-perturbation gene expression.

### 4.5 Graph attention network learns meaningful representations

To assess the effectiveness of the Graph Attention Network (GAT) in discovering meaningful patterns from the data, we analyze genes with the highest attention scores derived from the acquired attention maps after training. The GAT assigns attention values to each edge in the graph by aggregating the outgoing edges of each node (gene) within the network, typically using the sum aggregation method. Subsequently, genes are ranked based on these aggregated attention scores. To streamline the analysis, we focused on examining the attention weights of cell-type graphs in the Kang dataset post-training. This dataset involves a single-perturbation, interferon-beta (IFN-*β*), known for its potent antiviral activity and roles in mediating antiviral responses, inhibiting growth, and modulating immune functions [22]. Our findings indicate that the GAT identifies distinct gene sets across different cell types. Moreover, analysis of these high-attention genes (HAGs) suggests that such an approach can serve as an effective feature extraction method akin to differential gene expression analysis. Therefore, we conducted a comparison between the union of the top 50 HAGs and the union of the top 100 DEGs across PBMC cell types. Figure 10a shows a Venn diagram indicating the relationship between the HAGs and DEGs sets. The diagram reveals that 47.2% of the genes are exclusive to the HAGs category, signifying a significant portion of genes with high attention that are not differentially expressed. Conversely, 45.1% of the genes are exclusive to the DEGs category, underscoring a substantial number of differentially expressed genes that do not receive high attention. Notably, there is an overlap of 7.7% between HAGs and DEGs, signifying a small subset of genes that are both highly attended to and differentially expressed. This overlap suggests that while there is some intersection between high attention and differential expression, the majority of genes fall into one category or the other, indicating distinct regulatory or functional roles within the biological system being studied.

**Figure 10:**
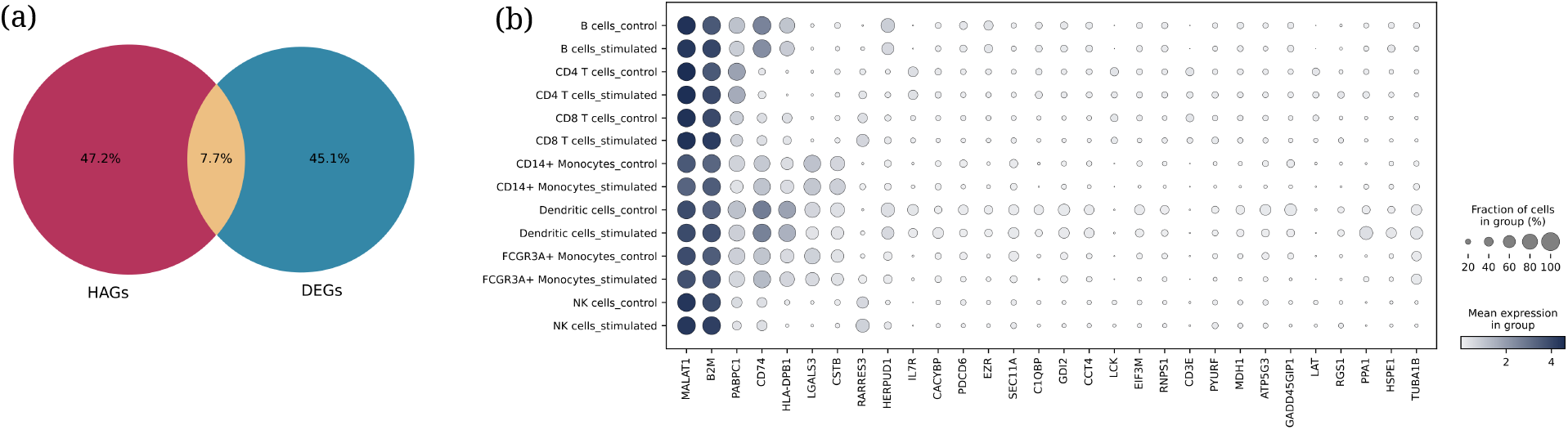
High attention gene analysis. (a) Venn diagram illustrating the relationship between the union of the top 50 HAGs and the union of the top 100 DEGs across PBMC cell types. (b) The gene expression state of the top 30 highly expressed genes from the HAGs that do not overlap with the DEGs set.

To this end, we investigate the set of genes from the HAGs that do not overlap with the DEGs to uncover meaningful biological patterns. Figure 10 illustrates the gene expression profiles of the top 30 highly expressed genes from this set. We further explore relevant experimental studies in the literature to identify associations between these genes and the perturbations or cell types in which they are highly expressed. For example, **CD74** is one of the top 20 HAGs in CD14+ monocytes and plays a critical role in antigen presentation and immune regulation within the immune system [27]. Another study by Nguyen-Pham et al. [28] demonstrated that CD74 negatively regulates dendritic cell migration, highlighting its importance in designing protocols to enhance the migratory ability of dendritic cells in immunological studies. **CD74, HLA-DQA1**, and **B2M** are HAGs in CD14+ monocytes and are involved in the antigen processing and presentation pathway, according to the KEGG pathways database. **LCK, LAT**, and **CD3E** are among the top HAGs in CD4 T cells and are members of immune system pathways such as T-cell receptor signaling and Th17 cell differentiation. See Figure 10. These genes serve as examples of cell type markers identified through our analysis.

To identify associations between interferon perturbation and Highly Associated Genes (HAGs), we conducted pathway enrichment analysis using the `prerank` function from the GSEApy package [29] in Python. This process involved ranking genes based on their attention scores and identifying the top HAGs involved in interferon-related pathways. For example, **POLR3G** and **POLR2K** are implicated in the positive regulation of type I interferon production (GO:0032481) and regulation of type I interferon production (GO:0032479) pathways. Additionally, we performed the same analysis for the overlap set between HAGs and DEGs, finding that **XAF1** is involved in multiple interferon-related pathways, including cellular response to type I interferon (GO:0071357) and type I interferon signaling pathway (GO:0060337). Furthermore, **XAF1** is identified as a uniform IFN-*β* marker according to [30].

Despite the Graph Attention Network’s (GAT) ability to attend to meaningful biological features, we investigate its capability to learn the correlation structures defining the edges in cell-type-specific graphs. To explore this, we compute Spearman’s correlation between the number of neighbors and the attention rank for the genes in each cell type. The results revealed a minimum correlation value of 0.60 observed in CD4T cells and a maximum value of 0.85 in FCGR3A+ monocytes. These correlation values across different cell types suggest that the GAT effectively learns the correlation network structure within cell-type-specific graphs.

## 5 Conclusion

In this study, we leverage a network representation of various cell types as an inductive bias to predict cell-type-specific responses to chemical perturbations. We demonstrated the effectiveness of our approach in predicting the mean and variance of post-perturbation expression for the DEGs across diverse cellular contexts. Notably, we tested our method on the most challenging cell types to ensure its generalizability in out-of-distribution scenarios. By employing the graph attention network algorithm, we extracted meaningful features from the cell-type-specific networks and compared this approach with standard differential gene expression analysis. Our comparative analysis with state-of-the-art generative models underscores the advantages of incorporating cell-type-specific information in predicting post-perturbation gene expression. The results confirm that our method performs well even in the most difficult cellular contexts, highlighting its potential for broad application in single-cell data analysis.

## 6 Data and Code Availability

All the experiments and datasets in this study are available in Github: https://github.com/reem12345/Cell-Type-Specific-Graphs

## Supplementary Figures

**Figure S1:**
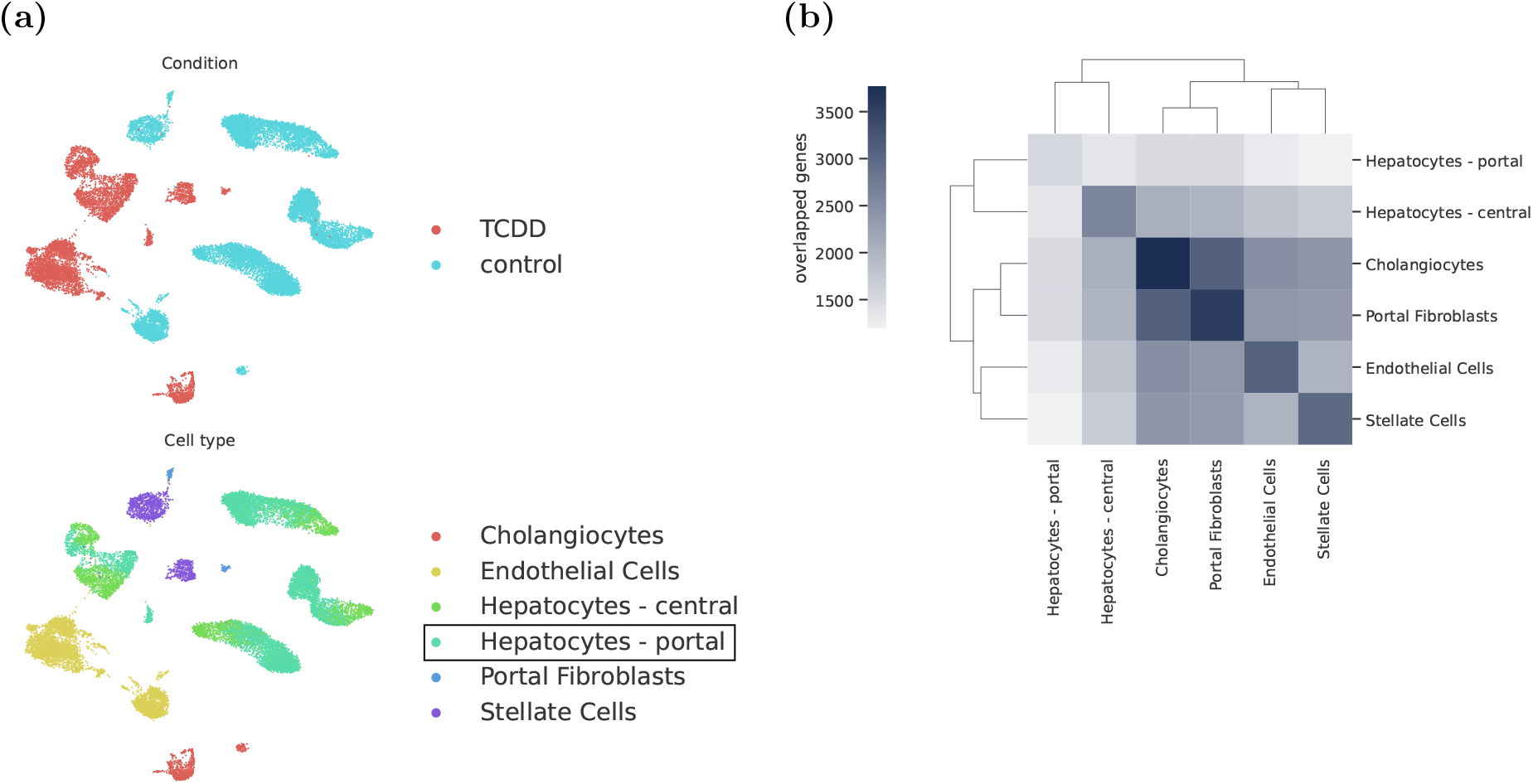
Single-perturbation experiment with Nault dataset. (a) UMAP Projection showing the testing cell type. (b) A clustered heatmap showing the number of shared genes/nodes between the cell-type graph pairs.

**Figure S2:**
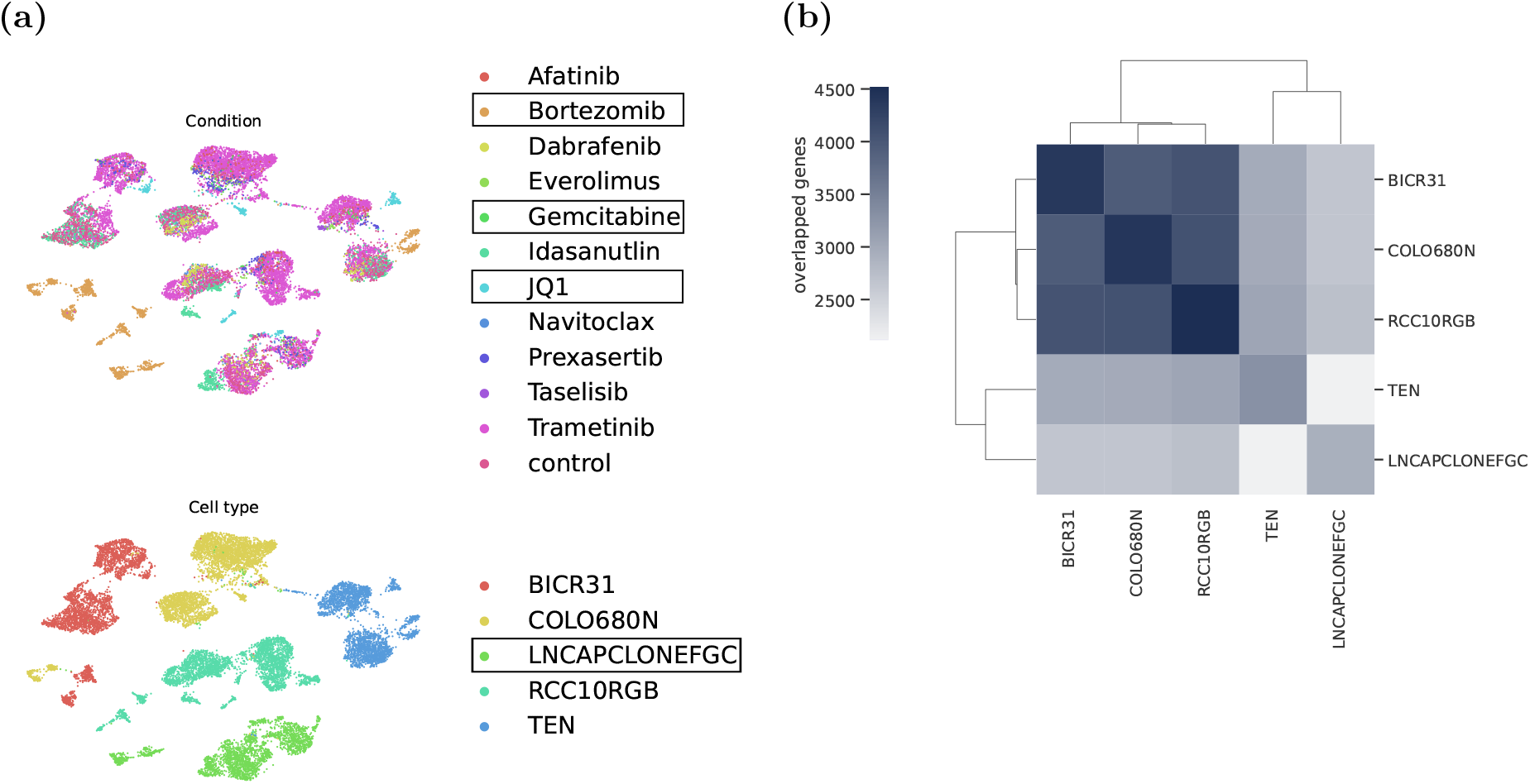
Multi-perturbation experiment with McFarland dataset. (a) UMAP Projection showing the testing cell type and drugs. (b) A clustered heatmap showing the number of shared genes/nodes between the cell-type graph pairs.

